# Grandparent inference from genetic data: The potential for parentage-based tagging programs to identify offspring of hatchery strays

**DOI:** 10.1101/2021.08.11.455946

**Authors:** Thomas A. Delomas, Matthew Campbell

## Abstract

Fisheries managers routinely use hatcheries to increase angling opportunity. Many hatcheries operate as segregated programs where hatchery-origin fish are not intended to spawn with natural-origin conspecifics in order to prevent potential negative effects on the natural-origin population. Currently available techniques to monitor the frequency with which hatchery-origin strays successfully spawn in the wild rely on either genetic differentiation between the hatchery- and natural-origin fish or extensive sampling of fish on the spawning grounds. We present a method to infer grandparent-grandchild trios using only genotypes from two putative grandparents and one putative grandchild. We developed estimators of false positive and false negative error rates and showed that genetic panels containing 500 - 700 single nucleotide polymorphisms or 200 - 300 microhaplotypes are expected to allow application of this technique for monitoring segregated hatchery programs. We discuss the ease with which this technique can be implemented by pre-existing parentage-based tagging programs and provide an R package that applies the method.

## [A] Introduction

Fisheries managers have long used hatcheries to increase angling opportunity and to compensate for anthropogenic impacts that have decreased population sizes (Waples et al. 2007). In some situations, it has been observed that hatchery-origin fish have lower fitness in the wild relative to natural-origin conspecifics, potentially due to selection in the hatchery environment or the use of hatchery strains that are not locally adapted (Ford 2002; Miller et al. 2004; Araki et al. 2007a, 2007b, 2008; Christie et al. 2014). In order to prevent negative effects of hatchery- and natural-origin fish interbreeding, it has been recommended that hatcheries operate as either integrated or segregated programs (Hatchery Scientific Review Group 2009). Integrated programs aim to balance the proportion of natural-origin fish in the hatchery broodstock and the proportion of hatchery-origin fish spawning naturally to minimize the effect of domestication (Goodman 2005). Segregated programs intend to minimize gene flow between hatchery- and natural-origin populations (Mobrand et al. 2005; Hatchery Scientific Review Group 2009). These strategies, developed in the context of Pacific salmon hatcheries, represent alternative approaches to both protect natural-origin populations and provide harvest opportunities for anglers.

In order to evaluate the efficacy of segregated hatchery programs, a method of monitoring gene flow from hatchery-origin fish to nearby natural-origin populations is needed. This has been previously estimated in several ways. Some studies estimate the proportion of fish on the spawning grounds that are of hatchery origin (pHOS) through observation of marks and/or tags present on hatchery fish (Dauer et al. 2009; Tattam and Ruzycki 2020). This metric is important as it demonstrates the potential for competition between hatchery- and natural-origin fish on the spawning grounds and suggests that hatchery introgression may be occurring. Alternatively, if the hatchery- and natural-origin populations show sufficient genetic differentiation, migration rates can be estimated from temporal samples (van Doornik et al. 2013), or genetic structure of hatchery- and natural-origin populations can be evaluated to determine whether the pattern indicates hatchery introgression (Matala et al. 2012; Ozerov et al. 2016; Lehnert et al. 2020). All of these approaches have drawbacks. Observing the proportion of hatchery-origin fish on the spawning grounds, while suggestive, does not directly assess gene flow as the reproductive success of hatchery-origin fish is unknown. Methods utilizing genetic differentiation are not applicable to cases without sufficient differentiation between the hatchery- and natural-origin populations, which is common where hatchery stocks were derived from nearby natural-origin populations.

An alternative technique uses genetic samples to infer relationships between hatchery broodstock and individuals sampled in the wild. Hatchery broodstock can be genetically sampled at the time of spawning, and their genotypes later used to infer whether a given fish is a descendent of hatchery broodstock. Parentage-based tagging (PBT) uses this approach to identify offspring of the hatchery broodstock for monitoring and management of hatchery stocks (Anderson and Garza 2005, 2006). Parentage-based tagging has been implemented and validated on large and small scales for a variety of species (DeHaan et al. 2008; Denson et al. 2012; Evans et al. 2018; Bingham et al. 2018; Campbell et al. 2019; Vandeputte et al. 2021), most notably Pacific salmonids (Steele et al. 2013, 2019; Beacham et al. 2019).

With appropriate methods for statistical inference, the general approach of PBT can be extended to identify grandchildren of hatchery broodstock. Genetic samples can be taken from hatchery broodstock, and samples from natural-origin fish can later be assessed to determine if they are grandchildren of those broodstock (and therefore had an unsampled, hatchery-origin parent). The relationship being inferred is a grandparent-grandchild trio consisting of one grandchild and two grandparents on the same side (i.e., either both maternal or both paternal grandparents). The other two grandparents and the parents are unsampled and therefore have unknown genotypes (Figure 1). While comprehensive sampling in the hatchery is straightforward, similar sampling of adults spawning naturally is often logistically prohibitive. The ability to infer recent hatchery ancestry without sampling naturally spawning parents would overcome this issue.

**Figure 1.**
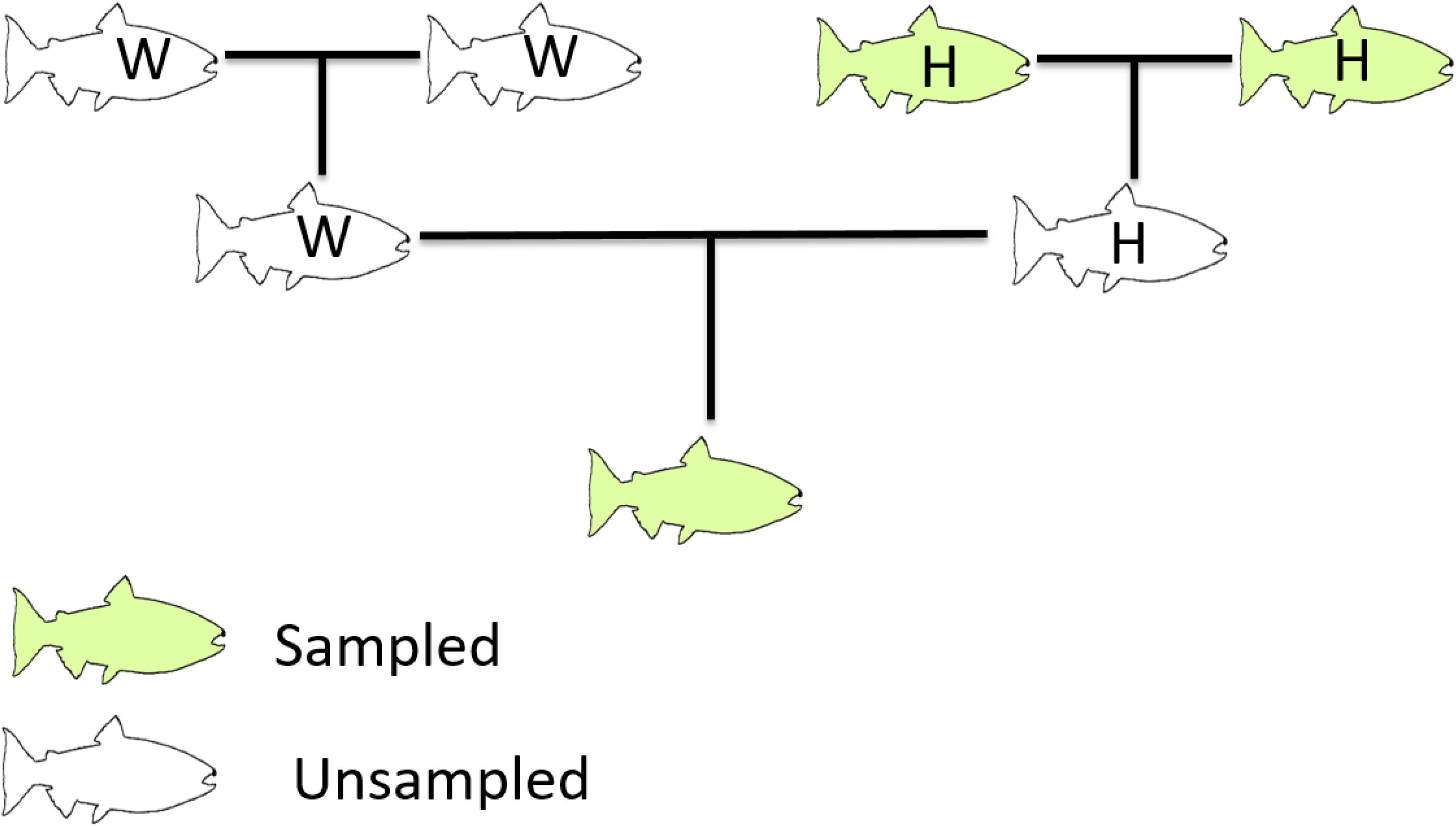
The pedigree the proposed method could be used to infer. H: hatchery-origin, W: natural-origin

Methods have previously been developed for inferring grandparent-grandchild relationships, but these methods are not optimal for inferring grandparent-grandchild trios. Letcher and King (2001) describe a method for inferring the relationship between all four grandparents and a grandchild. To identify offspring of hatchery strays, this method requires sampling natural-origin grandparents as well as hatchery broodstock which is not feasible in many situations. Christie et al. (2011) describe a method to infer grandparent-grandchild trios by identifying trios with no observed Mendelian incompatibilities (or less than a specified number). While this method has been successfully applied to situations where one parent is known (Christie et al. 2011; Sard et al. 2016), the lower power of exclusionary methods compared to likelihood-based methods (Anderson and Garza 2006) make exclusionary methods less feasible when no parents are sampled.

We here report a likelihood-based method to identify grandparent-grandchild trios using genotypes. Additionally, we develop techniques to estimate assignment error rates for this method and provide an R package implementing the described techniques (https://github.com/delomast/gRandma).

## [A] Methods

### [C] Inferring grandparent-grandchild relationships

Relationships have previously been inferred from genetic data using likelihood ratios to compare the putative relationship (e.g. parent-offspring) to an alternative relationship (typically that the individuals are unrelated) (Marshall et al. 1998; Anderson and Garza 2006; Kalinowski et al. 2007; Anderson 2012). Methods for calculating likelihoods of relationships (simple pedigrees) have previously been detailed (Thompson 1976, 2000; SanCristobal and Chevalet 1997; Anderson and Garza 2006), and we extended this approach specifically to grandparent-grandchild trios. Given allele frequencies for a locus under Hardy-Weinberg equilibrium (HWE), the likelihood of the three individuals being unrelated was calculated as the product of the probability of sampling each genotype from the population. The likelihood of the individuals being a grandparent-grandchild trio was calculated by utilizing allele frequencies and the laws of Mendelian inheritance. Genotyping error was accounted for by marginalizing the true genotypes utilizing estimates of genotyping error (Anderson and Garza 2006). This allows genotyping error to be flexibly modeled in any way that yields, for a given true genotype, the probability of observing each genotype. With the R package we provide, users can specify these probabilities or utilize a default error model. The default error model, used in the analyses described in this study, is described in supplementary file 1. Loci were considered to be independent, as in previous methods of relationship inference and pedigree reconstruction (Marshall et al. 1998; Riester et al. 2009; Jones and Wang 2010; Anderson 2012; Huisman 2017).

Calculating likelihoods for large numbers of possible trios can be computationally intensive. We therefore implemented a preliminary screen based on the observed number of Mendelian incompatibilities (MI) in a grandparent-grandchild trio. Trios with more than *m* MIs were excluded from consideration. The value of *m* was chosen so that the probability of excluding a true grandparent-grandchild trio was less than 0.0001 for a trio with no missing genotypes. With a constant value of *m*, the probability of rejecting a trio with one or more missing genotypes is smaller, as a locus with a missing genotype cannot be considered a Mendelian incompatibility. The probability of rejecting a true trio given a value of *m* was calculated by representing the number of MIs as a Markov Chain, following the method detailed by Anderson (2012) but extended to the relationship of grandparent-grandchild trio.

Assigning grandparent-grandchild relationships can be treated as a hypothesis test by comparing the calculated log-likelihood ratio (LLR) to a critical value, *c* (SanCristobal and Chevalet 1997; Anderson and Garza 2006). If the LLR was greater than or equal to *c*, the trio was considered related; otherwise, the trio was considered unrelated. The value of *c* can be chosen to achieve a desired balance of false negative and false positive error rates.

### [C] False positive error rates

Per-comparison false positive error rates (probability of assigning a relationship to a trio that is not a grandparent-grandchild trio) are dependent upon the true relationship for a trio. We have implemented methods to assess false positive error rates for unrelated trios (typically the most important) as well as 13 other types of relationships. In these relationships, the two putative grandparents were considered unrelated to each other, but the putative grandparents had different relationships to the putative grandchild. The relationships considered represented combinations of true grandparents, individuals unrelated to the putative grandchild, great-aunts, half-great-aunts, and first cousins of the putative grandchild”s true grandparent.

Estimating the per-comparison false positive error rate for a given value of *c* has been demonstrated for parent-offspring and sibling relationships using importance sampling (Anderson and Garza 2006; Baetscher et al. 2018), a Monte Carlo variance reduction technique. Variance reduction is needed as a naïve Monte Carlo approach would be inefficient at estimating very small false positive rates (Anderson and Garza 2006). Small error rates can be meaningful as the experiment-wide false positive error rate is estimated by the product of the per-comparison false positive error rate and the number of trios (with the corresponding true relationship) considered. We extended this approach to the current application of assessing grandparent-grandchild trios. We also implemented an alternative method utilizing stratified sampling, another Monte Carlo variance reduction technique, because specific implementations of importance sampling can produce unreliable results (Owen 2013).

### [C] Importance sampling

A standard Monte Carlo estimator of false positive rates would be obtained by simulating genotypes for trios of a given relationship (such as unrelated) and recording how many fit the criteria to be considered related. Importance sampling can be thought of as focusing the simulation on producing mostly genotypes that do assign and then correcting for this modification. To implement importance sampling, we simulated genotypes from the distribution of genotypes in true grandparent-grandchild trios. Missing genotypes were accounted for utilizing the forward-backward algorithm described below with the state of the Markov chain representing the number of missing genotypes in one individual. If the simulated trio had fewer than *m* MIs and the calculated LLR was greater than or equal to *c*, the observation was recorded as a false positive with the appropriate importance sampling weight (Owen 2013).

### [C] Stratified sampling

Similar to importance sampling, the goal of stratified sampling was to focus simulation effort on categories (strata) that produce false positives. We stratified the distribution of trio genotypes by the number of observed MIs. This was a natural choice because the algorithm we use explicitly filters possibilities based on the number of observed MIs. Therefore, we can eliminate simulating genotypes for most strata. Sampling effort can then be focused on trios with *m* or fewer MIs. This method requires calculating the probability that a trio of given relationship has a given number of observed MIs and the ability to simulate genotypes for a trio given a relationship and number of MIs. Genotypes for trios were simulated in each stratum and the false positive rates were recorded. Utilizing the probabilities that a trio has each number of MIs (i.e., the size of the strata), the overall false positive rate was then calculated.

To calculate the probability a trio has a given number of observed MIs, we represented the observation of MIs and missing genotypes as a Markov chain and utilized the forward step of the forward-backward algorithm. We extended the approach described by Anderson (2012) to account for the common practice of only analyzing samples given a maximum number of missing genotypes, *d*, and to fit the target relationships. The value of *d* in the current analyses was 10% of loci. Let *s*_*i*_ be the state, describing the number of observed MIs and missing genotypes, after observing locus *i*. Prior to observing any loci, *s*_0_ = (0,0,0,0), representing the number of observed incompatibilities and number of missing genotypes for the three individuals. Let *a*_*i*_ be a vector indicating whether an incompatibility or any missing genotypes are observed at locus *i*, in the same order as *s*_*i*_. Assuming HWE, known allele frequencies, known locus-specific probabilities of a genotype being missing, and observation of missing genotypes is independent across loci and individuals, the probabilities of each possibility for *a*_*i*_ can be calculated according to standard probability arguments for a given true relationship. The probability of being in state *x* after a given locus can then be calculated as

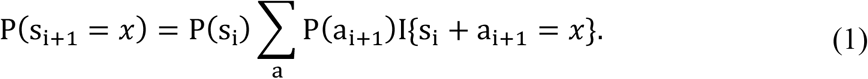

This can be evaluated recursively to obtain the probabilities of each final state. The probability a trio has a given number of observed MIs conditional on *d* can then be calculated.

However, memory constraints make saving all the probabilities at each step impractical even with moderate numbers of loci. We can save only the probabilities of states with *m* or fewer MIs and with all three individuals having *d* or fewer missing genotypes to reduce memory usage. These values are divided by the probability that all members have *d* or fewer missing genotypes to obtain probabilities of being in each state conditional on *d*. The probability of observing an individual with more than *d* missing genotypes can be obtained through the same algorithm (forwards step), but with *s* and *a* now only representing missing genotypes in one individual. Because we assumed that missing genotypes are independent between individuals (given locus-specific rates of missing genotypes), the probability of all three samples having *d* or fewer missing genotypes is straightforward to calculate using the obtained probability that one individual has more than *d* missing genotypes.

To utilize stratified sampling, we need to simulate genotypes for trios with a specified number of MIs. The backwards step of the forward-backward algorithm fills this need. Given *L* loci, a value of *s*_*L*_ is chosen given the number of MIs by sampling a categorical distribution with probabilities proportional to the probability of each *s*_*L*_ that has the specified number of MIs (and allowable number of missing genotypes). Next, for each locus and iterating backwards, a value for *a*_*i*_ is chosen by sampling a categorical distribution with

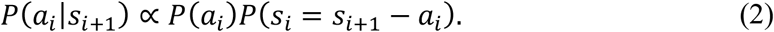

Once *a*_*i*_ is chosen, genotypes are sampled using the genotype frequencies for the true relationship calculated from the allele frequencies, HWE, laws of Mendelian inheritance, and genotyping error rates. If all three genotypes are observed (i.e., no missing genotypes in the chosen *a*_*i*_), then the genotypes are either sampled conditional upon an MI being present or not.

### [C] False negative error rate

To estimate the per-comparison false negative error rate (probability of failing to assign a relationship to a true grandparent-grandchild trio) for a given value of *c*, Monte Carlo methods have been previously used for other relationships (Anderson and Garza 2006; Baetscher et al. 2018) and we adopted this strategy here. Genotypes of grandparent-grandchild trios are simulated and loci with missing genotypes for each individual are chosen by representing the observation of missing genotypes as a Markov chain and utilizing the forward-backward algorithm, as described above with the state representing the number of missing genotypes in one individual. Log-likelihood ratios were calculated for the simulated genotypes, and the proportion of trios with the number of MIs greater than *m* or LLR less than *c* was the estimate of the false negative rate.

### [C] Panel size simulations and error rate estimator evaluation

A key question for designing experiments implementing this method is, how many loci need to be genotyped to obtain reliable assignments? To help answer this question, we estimated error rates for panels of different size. We simulated panels containing 100, 300, 500, 700, and 900 biallelic single nucleotide polymorphisms (SNPs) and panels containing 100, 200, 300, and 400 triallelic microhaplotypes. The SNPs and microhaplotypes were simulated with expected heterozygosity of 0.22 (allele frequencies of 0.125 and 0.872) and 0.42 (allele frequencies of 0.13, 0.13, and 0.74), respectively. This choice reflects the mean expected heterozygosities for SNPs and microhaplotypes in a comparison of both marker types for relationship inference in rockfish (Baetscher et al. 2018). The power of a given marker for kinship inference is directly related to its variability (Anderson and Garza, 2006). While we investigated marker panels of differing size but fixed variability, similar power to the simulated panels could be achieved with fewer, more variable markers. The probability of a missing genotype at a locus was set at 3% in these analyses. It is common practice to remove any samples with more than a threshold number of missing genotypes, and so we restricted all simulated genotypes to have 90% or more genotypes present. Given a true relationship of unrelated, false negative and false positive error rates were estimated for a range of *c* values and the relationship between error rates was compared between panels. Integer values of *c* were chosen starting at 0 and increasing until the estimated false negative rate was above 0.05. Estimates of false negative rates and estimates from the importance sampling method were derived from 10,000 and 1,000,000 Monte Carlo iterations, respectively. Estimates from the stratified sampling routine were derived from 1,000,000 iterations for each stratum (number of observed Mendelian incompatibilities) less than or equal to *m*.

To compare performance of the two methods for estimating false positive error rates, we compared the estimated error rates for unrelated trios between the methods with all four simulated microhaplotype panels. Estimates for other true relationships were compared using the 300 locus microhaplotype panel. Finally, to examine the importance of modelling the presence of missing genotypes, we compared importance sampling estimates using the 300 locus microhaplotype panel with the probability of a genotype being missing equal to 3% and 0%.

The scripts used to perform these simulations and their outputs are available at https://github.com/delomast/gpError2021.

### [C] Example analyses

To fully evaluate false positive per-comparison error rates, one needs to have a general idea of the size of the analysis (number of comparisons) being attempted. We consider two examples modeled around steelhead *Oncorhynchus mykiss* hatchery programs in the Snake River basin using data collected during 2018. The Upper Salmon B-run (USB) represents a smaller hatchery program and spawned 66 steelhead in 2018. The Dworshak National Fish Hatchery (DNFH) represents a larger hatchery program and spawned 1,778 steelhead in 2018. Both of these hatchery programs take genetic samples from all broodstock and record the day of spawning, phenotypic sex, and crosses made. Data collected at the hatchery (or a genetic sex marker) can be used to constrain the number of possible pairs of grandparents considered in an analysis. For each hatchery program, we consider the effect on the desired per-comparison false positive rate of using no data, phenotypic sex, spawn day, phenotypic sex and spawn day, or cross records.

In the example analysis, we assume that natural-origin juveniles are sampled and that exact age is unknown but is constrained to one, two, or three years old. The effect of this assumption is that three potential years of parents must be considered for each juvenile. Hatchery-origin steelhead in the Snake River basin return almost exclusively as three and four year old fish (Warren et al. 2017), so this translates to four years of potential grandparents that must be considered.

The total number of comparisons in these analyses is the product of the number of possible pairs of grandparents per year, the number of years of potential grandparents being considered (four), and the number of potential grandchildren evaluated (assumed here to be 200). We then calculate the desired per-comparison false positive (true relationship of unrelated) error rate to achieve an expected number of false positive assignments of 0.1 (0.1 / number of comparisons) assuming all trios are unrelated. This ignores false positives arising from trios of other relationships. In some situations, error rates for alternative relationships are important to consider, but in analyses of segregated hatchery programs (where no breeding of hatchery-origin and natural-origin fish is desired), false positives arising from trios of other relationships are not necessarily harmful as the purpose is to identify fish with recent hatchery-origin ancestry.

## [A] Results

Estimated error rates declined with increasing panel size, and the microhaplotype panels showed lower error rates than SNP panels of similar size (Figure 2). For example, false positive error rates (true relationship of unrelated) below 10^−10^ were achieved at false negative error rates below 0.05 with SNP panels containing 700 or more loci and microhaplotype panels containing 300 or more loci. False positive rates for related trios decreased with decreasing relatedness between individuals in the trio (Supplemental Figure 1).

**Figure 2.**
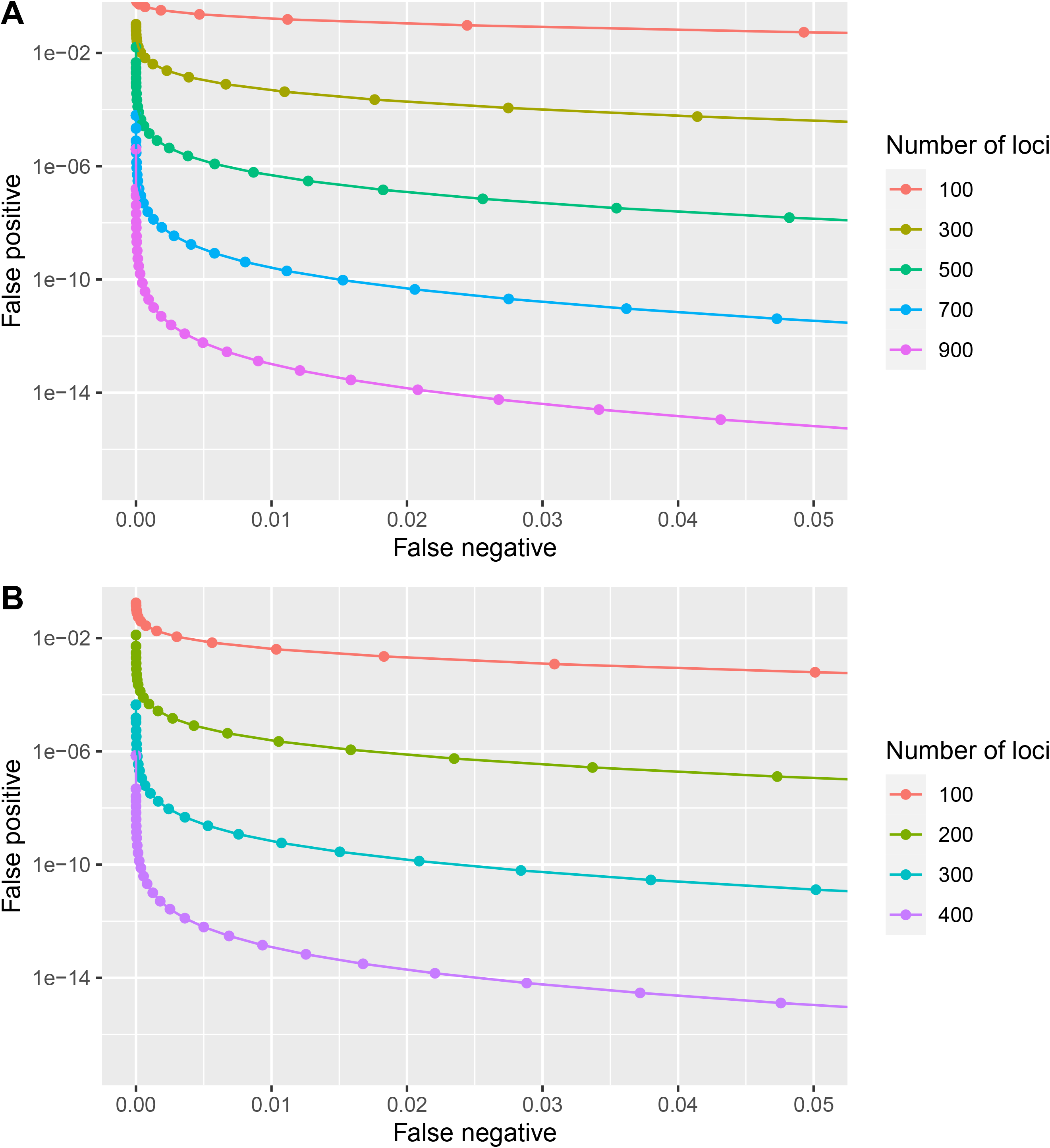
Error rates estimated by importance sampling for simulated (A) SNP and (B) microhaplotype panels with varying numbers of loci.

False positive rates estimated by importance sampling and stratified sampling were practically identical when estimates were above approximately 10^−8^ (Figure 3). As the false positive rate decreased below this, the importance sampling method estimated false positive rates higher than the stratified sampling method. In these cases, the stratified sampling method estimated false positive rates of 0 within one or more strata (i.e., no false positives were sampled out of the 1,000,000 iterations). Similar results were obtained for false positive rates estimated for trios with relationships other than unrelated (Figure 4) except for two trio types (true grandparent and unrelated; true grandparent and cousin of grandparent) that had noticeably different estimates between the two methods and did not have a low false positive rate.

**Figure 3.**
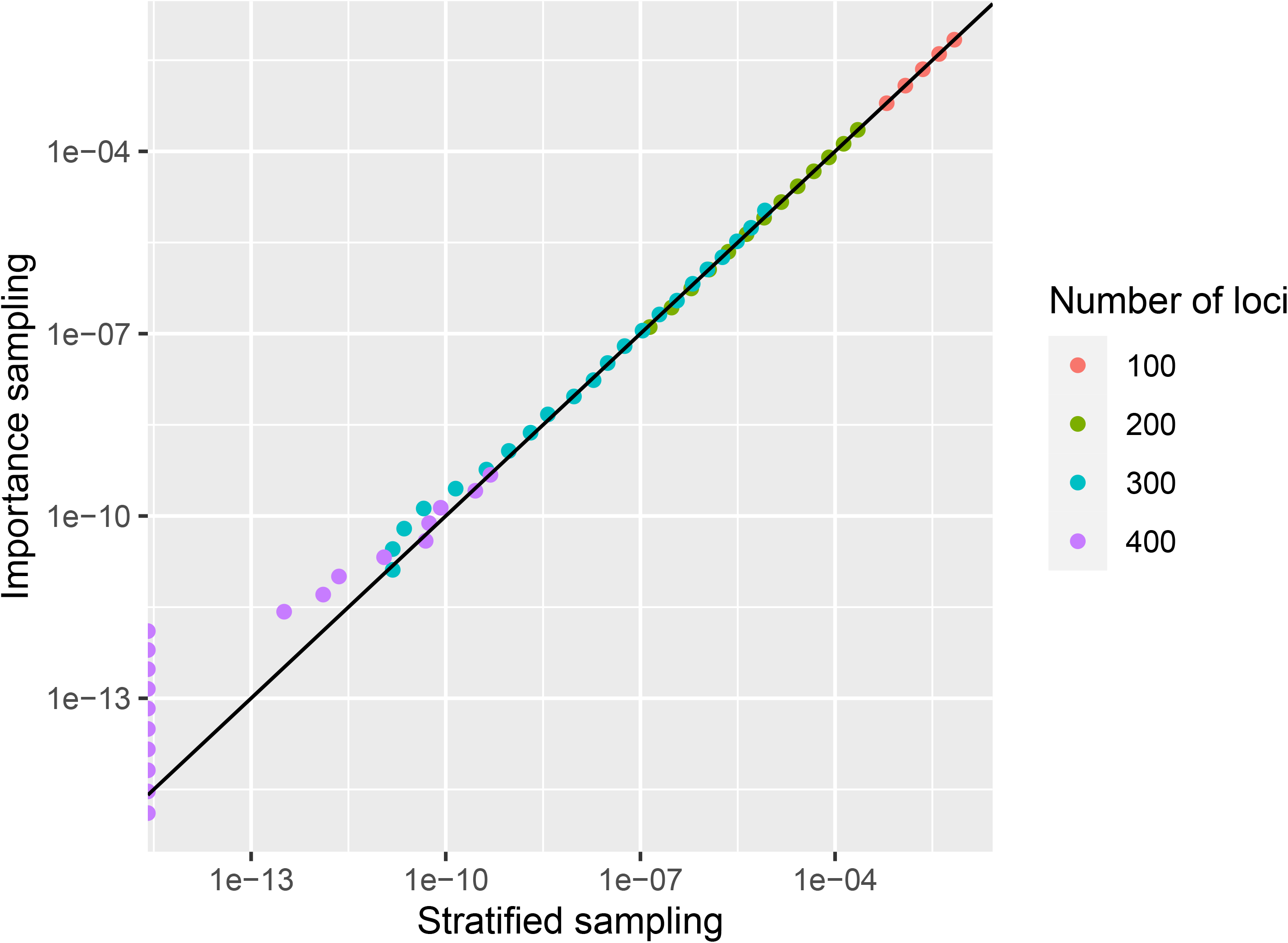
Comparison of false positive (true relationship of unrelated) error rates for simulated microhaplotype panels estimated by importance sampling and stratified sampling. The black line represents y = x. Points shown on the y-axis had an estimated error rate of 0 from stratified sampling.

**Figure 4.**
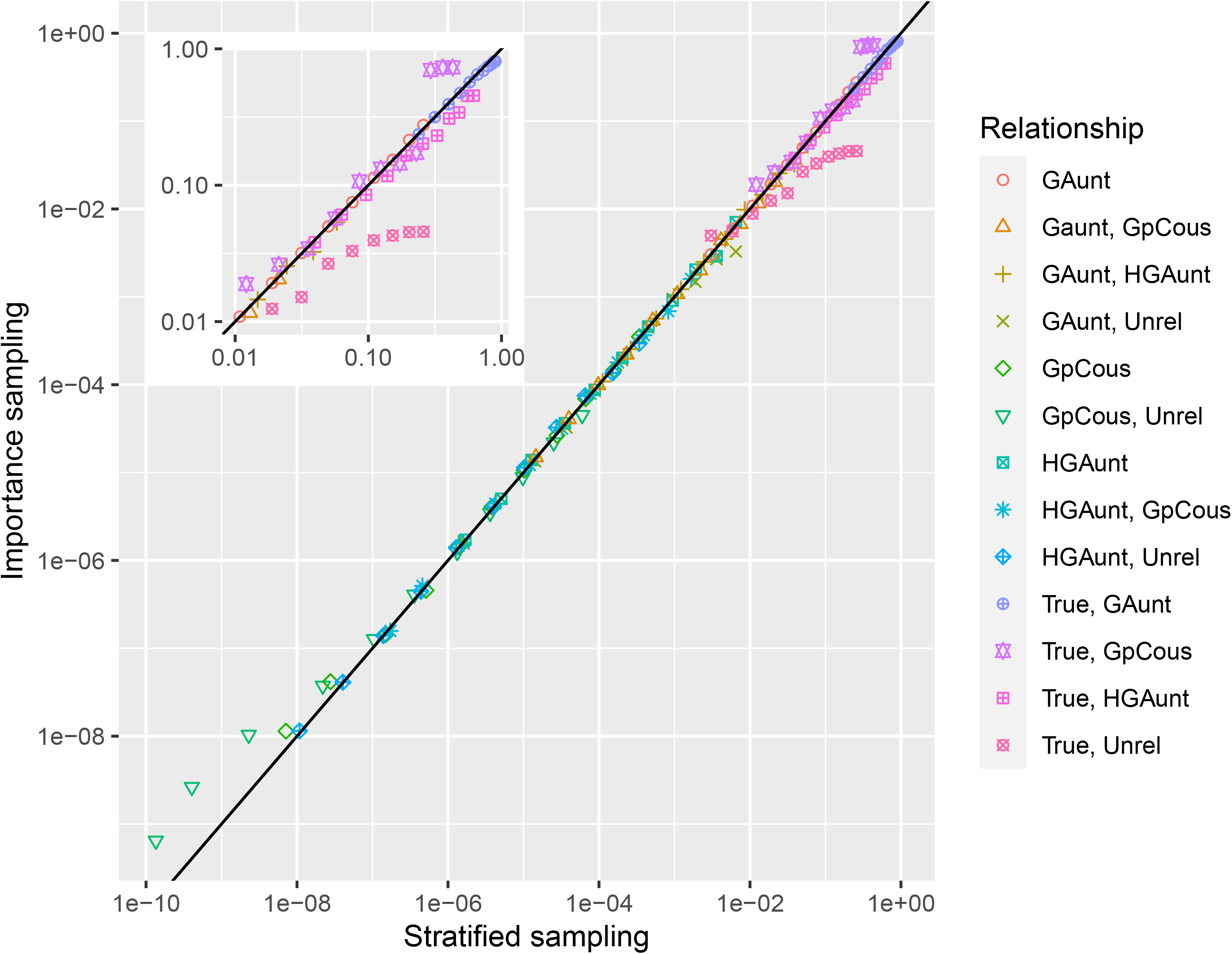
Comparison of false positive error rates estimated by importance sampling and stratified sampling for related trios with the simulated 300 locus microhaplotype panel. The black line represents y = x. The two putative grandparents were unrelated to each other. The relationship labels indicate the relationship of the two putative grandparents to the putative grandchild. Where the label is only one relationship, both putative grandparents have the same relationship to the putative grandchild. True is true grandparent, GAunt is great-aunt, HGAunt is half great-aunt, GpCous is first-cousin to the putative grandchild’s true grandparent, Unrel is unrelated. The inset gives a magnified view of the region containing error rates close to 1.

Incorporating a 3% missing genotype rate in the error rate estimation had a moderate effect on the results (Figure 5). For example, at a false negative rate of 0.05, the false positive (true relationship of unrelated) error rates were approximately (derived from linear interpolation with neighboring points) 1.3 . 10^−11^ and 2.1 . 10^−12^ when missing genotypes had rates of 3% and 0%, respectively.

**Figure 5.**
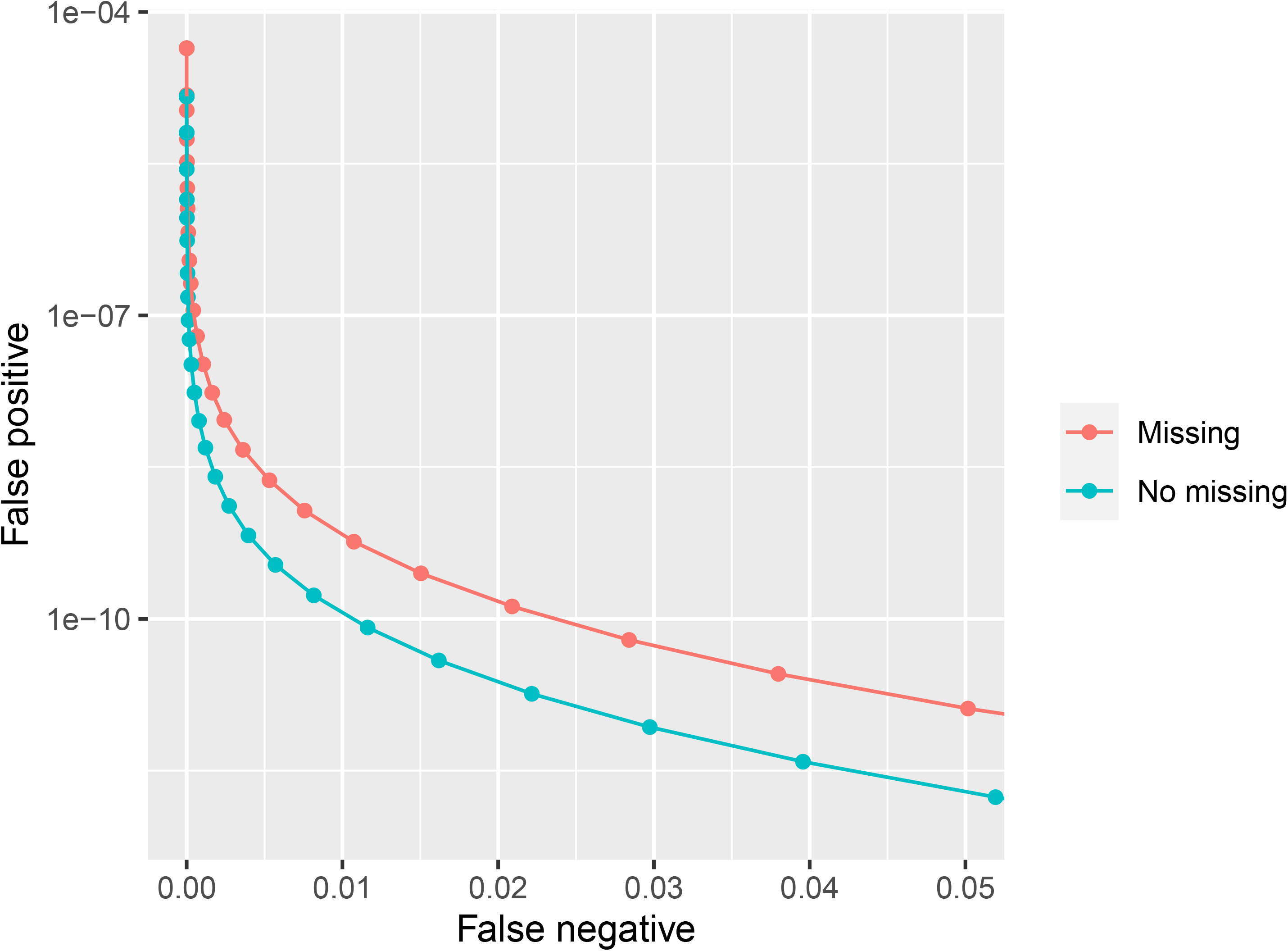
Error rates estimated by importance sampling for the simulated 300 locus microhaplotype panel with two rates of missing genotypes: 3% (Missing) and 0% (No missing). False positive error rate is the rate for unrelated trios.

The example hatchery programs considered show that analysis of smaller programs (e.g., USB) require per-comparison false positive rates on the order of 10^−6^ - 10^−8^ depending on what data, if any, are available to reduce the number of comparisons (Table 1). Larger programs, similar to DNFH, require rates on the order of 10^−7^ - 10^−10^ if the number of comparisons can be reduced by one of the data sources considered. Error rates estimated for the simulated panels (Figure 2) suggest that these rates can be achieved with panels containing 500 - 700 SNPs or 200 - 300 microhaplotypes. If no data are available to reduce the number of pairs of grandparents that must be considered, then hatchery programs similar in size to DNFH (1,700 fish spawned / year) will require error rates on the order of 10^−11^, which is achievable with panels containing 700 - 900 SNPs or 300 - 400 microhaplotypes.

**Table 1.** Desired per-comparison false positive (true relationship of unrelated) error rates for analyses using different data sources to reduce the number of comparisons.

| Hatchery | Data used | Number of<br>possible crosses<br>per year | Years of<br>potential<br>grandparents | Number of<br>potential<br>grandchildren | Number of<br>comparisons | Desired per-comparison false<br>positive error<br>rate |
| --- | --- | --- | --- | --- | --- | --- |
| USB | None | 4,290 | 4 | 200 | 3,432,000 | $2.91 \cdot 10^{-8}$ |
| | Sex | 1,088 | | | 870,400 | $1.15 \cdot 10^{-7}$ |
| | Spawn day | 584 | | | 467,200 | $2.14 \cdot 10^{-7}$ |
| | Spawn day and sex | 162 | | | 129,600 | $7.72 \cdot 10^{-7}$ |
| | Cross records | 34 | | | 27,200 | $3.68 \cdot 10^{-6}$ |
| DNFH | None | 3,134,670 | 4 | 200 | 2,507,736,000 | $3.99 \cdot 10^{-11}$ |
| | Sex | 769,348 | | | 615,478,400 | $1.62 \cdot 10^{-10}$ |
| | Spawn day | 284,270 | | | 227,416,000 | $4.40 \cdot 10^{-10}$ |
| | Spawn day and sex | 69,709 | | | 55,767,200 | $1.79 \cdot 10^{-9}$ |
| | Cross records | 1,003 | | | 802,400 | $1.25 \cdot 10^{-7}$ |

## [A] Discussion

The methods developed here facilitate direct monitoring of introgression between hatchery- and natural-origin populations without relying on genetic differentiation. This fills an unmet need for monitoring the segregated hatchery programs upon which many fisheries rely. The method identifies fish with an unsampled, hatchery-origin parent using genetic samples from the hatchery broodstock and estimates error rates given allele frequencies for a population.

Another benefit of this method is the ease with which it can be implemented through existing PBT programs because the hatchery and laboratory processes required to sample and genotype broodstock are already in place. Additional requirements for grandparent inference would be the development of a genetic panel with sufficient power (given the size of the relevant hatcheries) and collection of samples from natural-origin fish that are potential grandchildren of the hatchery broodstock. In the past decade, there have been multiple, large-scale demonstrations of the efficacy of PBT for monitoring fisheries (Steele et al. 2013, 2019; Beacham et al. 2019). Numerous PBT programs have been recently reported (Evans et al. 2018; Bingham et al. 2018; Campbell et al. 2019; Vandeputte et al. 2021) and current tagging technologies can be replaced by PBT to provide additional benefits at similar or reduced costs (Beacham 2021). This implies that PBT will continue to grow in usage, and the method described here will become feasible for a larger number of hatchery programs.

The simulated microhaplotype panels demonstrated that error rates low enough for relatively large analyses of segregated hatchery programs are achievable with panels containing 300 - 400 microhaplotypes. The size/power required from the genetic marker panel will depend on the data available to limit the number of comparisons. As has been previously demonstrated, the availability of accurate hatchery cross-records greatly reduces the power required for grandparent inference (Letcher and King 2001), but we note that simply having sex information associated with a broodstock individual”s genotype can have a sizeable effect (Table 1). Panels genotyping hundreds of loci have been created using cost-effective, amplicon sequencing techniques (Campbell et al. 2015; Janowitz-Koch et al. 2019), and so we conclude that grandparent inference with this method is feasible using current genotyping techniques.

Application of this method at larger scales will require considering potential grandparents from multiple hatcheries. In these situations, having data available to reduce the number of comparisons can be critical for making an analysis feasible. For example, if natural-origin steelhead are sampled at Lower Granite Dam on the Snake River (Hargrove et al. 2021), then all steelhead hatcheries adjacent to and upstream of the dam must be considered (approximately 5,000 total steelhead spawned annually). With no data other than the year and hatchery at which fish were spawned, the number of potential pairs of grandparents would be approximately 5.5 . 10^6^ each year, while considering phenotypic (or genetic) sex would reduce this to 1.4 . 10^6^ (IDFG, unpublished).

We expect this method to be most applicable for assessing segregated hatchery programs. For some closely related trios, false positive error rates were relatively high with the 300 microhaplotype panel (Supplemental Figure 1). For all analyses, the impact of these error rates is mediated by the infrequency of comparisons involving closely related individuals. When analyzing segregated programs, false positives from closely related trios may not negatively impact conclusions as they indicate recent hatchery ancestry. However, when analyzing integrated hatchery programs, distinguishing trios with different relationships can be important. Application of this method could then either be infeasible or require a more powerful genetic panel. Additionally, the number of relationships for which we provide estimators covers trios with a range of relatedness but is not exhaustive. In some specific cases, other relationships may be present at impactful frequencies. For example, if generations overlap substantially then the impact of trios containing aunts, half-aunts, and first-cousins may need to be considered.

The method developed here addressed shortcomings of previously developed methods for grandparent inference with moderately sized genetic panels. Previous methods required either that all four grandparents were sampled (Letcher and King 2001) or used an exclusionary method that inherently had lower power than likelihood-based methods (Christie et al. 2011). Additionally, the exclusionary method did not provide a formal treatment of genotyping error. Applications of the exclusionary method (Christie et al. 2011; Sard et al. 2016) have utilized panels of tens of microsatellites. When using SNPs or microhaplotypes, hundreds or thousands of loci are typically genotyped and minimal error rates (e.g., 1%) still result in errors being present in a majority of individuals. An additional effect of ignoring genotyping error in an exclusionary method is that false negatives are implicitly assumed not to occur. It is worth noting that use of a Markov chain to model the number of observed Mendelian incompatibilities and missing genotypes in a trio, as we developed here, could be incorporated into the exclusionary method. This would allow both the formal incorporation of genotyping error and estimation of false positive and false negative error rates.

Importance sampling and stratified sampling have different weaknesses, making each better suited to different situations. For example, importance sampling can perform suboptimally when the estimate is dominated by a small fraction of samples (Owen 2013). Stratified sampling does not have this same drawback, but it can fail to give a meaningful estimate when the false positive rate within a particular stratum is low enough that a reasonable number of samples cannot estimate it accurately. Comparison of the importance sampling and stratified sampling methods showed that when the true relationship was unrelated, they estimated essentially the same false positive rate when that rate was above approximately 10^−8^. This suggests the importance sampling routine performed well for unrelated trios. At lower false positive rates, the stratified sampling estimates were lower (and in some cases 0) than those from importance sampling. In these cases, one or more strata had estimates of 0, indicating that the 1,000,000 samples taken were not sufficient to observe one or more false positives. This demonstrates one of the strengths of importance sampling compared to stratified sampling - some situations will have false positive rates small enough they cannot be efficiently estimated by this stratified sampling routine. This is further emphasized by the greater computational effort devoted to stratified sampling in this study (1,000,000 iterations in each stratum vs. 1,000,000 iterations total).

For false positive error rates under relationships other than unrelated, the importance sampling and stratified sampling estimates were again largely the same. In a few cases where one potential grandparent was the true grandparent and the other was not, the estimates were noticeably different and the stratified sampling method achieved at least 150 false positive observations in each strata. This suggests that for these relationships the importance sampling method may have performed suboptimally. Similar observations were made for importance sampling estimates of false positive error rates in parentage inference, where performance of a given importance distribution varied depending on the true relationship for which an error rate was estimated (Anderson and Garza 2006). One strategy would be to design an alternative importance distribution, but for closely related trios, error rates for most panels will likely be high enough that they are amenable to stratified sampling.

The current method assumes loci are in linkage equilibrium. If loci are physically linked (but still in equilibrium) the methods described here can be applied with some additional consideration. Physical linkage of two loci will result in grandchildren being more likely to inherit alleles at both loci from one of the two grandparents in a trio. The estimated false positive (when true relationship is unrelated) error rates are not affected because unrelated individuals are not impacted by physical linkage. For the other estimated error rates (false positives for trios with other relationships and false negatives), the simulated LLRs will have lower variance than the true distribution, causing error rates to be underestimated.

One drawback of the method as implemented is that computational efficiency decreases with increasing numbers of alleles per locus. This is partly due to the flexibility of the genotyping error model and the need to marginalize over all possible true genotypes. In the current implementation, computation is sped up by precomputing likelihood values for observed trio genotypes at each locus. This works well when the number of alleles per locus is small, but as the number of alleles increases, this can become impractically slow. As such, the current implementation will be most suitable to panels containing biallelic SNPs and microhaplotypes, which in our experience typically have three to five alleles. For panels of highly variable loci, a different implementation of this method, particularly with a more streamlined genotyping error model, would be necessary.

## [A] Acknowledgements

Funding for this project was provided by Bonneville Power Administration Project 2010-031-00. We would like to thank John Powell, John Hargrove, Matt Corsi, and Eric Anderson for providing comments that improved this manuscript. There is no conflict of interest declared in this article.

---

**Supplemental Figure 1.**
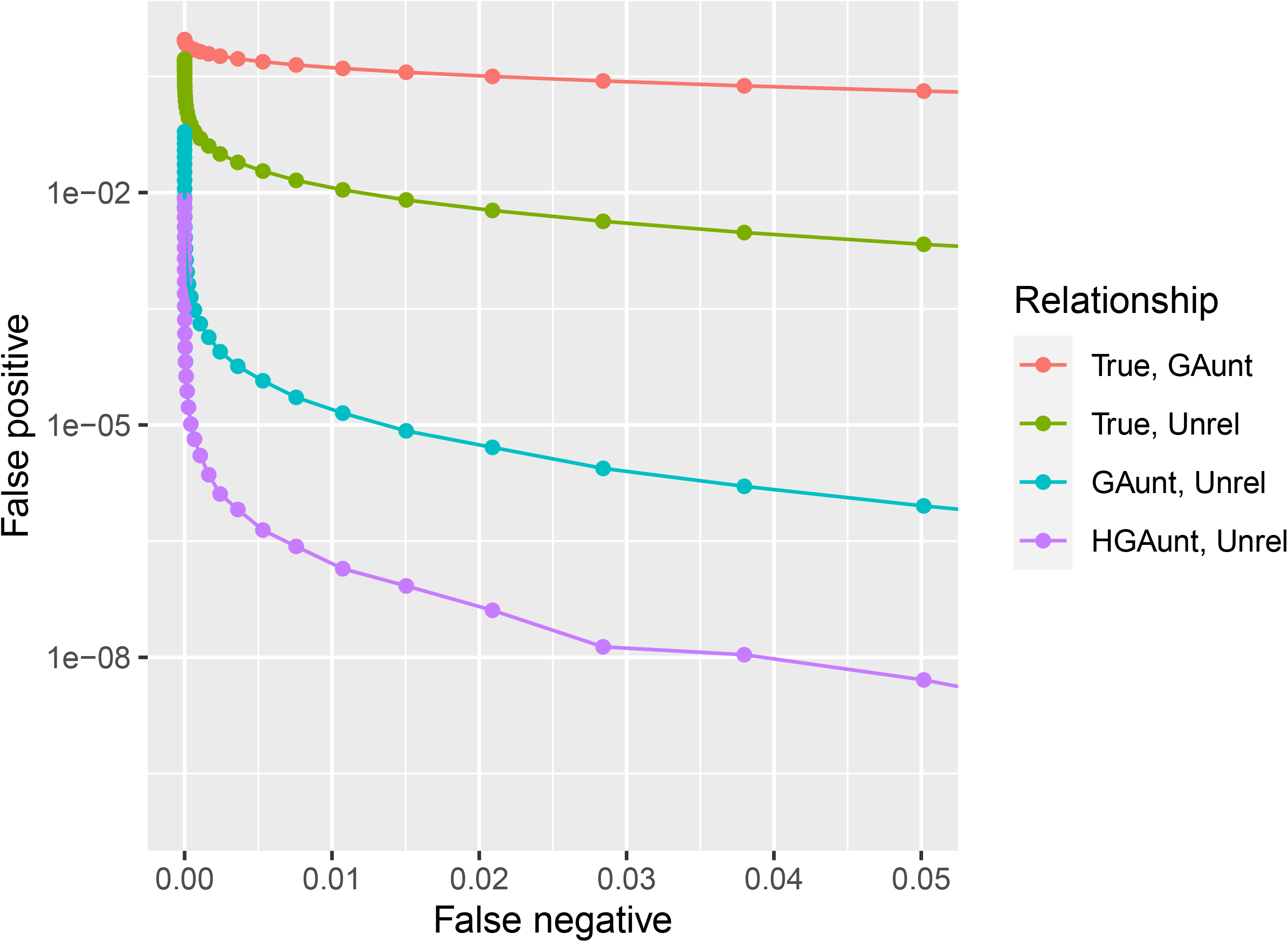
Error rates estimated by stratified sampling for the simulated 300 locus microhaplotype panel for select related trios. The two putative grandparents were unrelated to each other. The relationship labels indicate the relationship of the two putative grandparents to the putative grandchild. True is true grandparent, GAunt is great-aunt, HGAunt is half great-aunt, Unrel is unrelated.

## Supplementary file 1

### 1 Default error model for gRandma

Each locus is treated separately and each genotype is treated as two observations (one for each allele in the presumed diploid). For a given locus, the error model first relies upon a ”per allele error rate”, *ϵ*, which represent the probability that, when observing one allele, you observe any allele other than the true one. The default value is 0.005, or 0.5%. This value is used along with a measure of similarity between alleles to calculate the probability of observing each allele given the true allele. For a locus with *I* alleles, given the true allele *i*, the probability of correctly observing *i* is 1 *− ϵ* and the probability of observing allele *j* where *j* ≠ *i* is

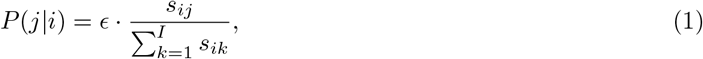

where *s*_*ij*_ is the similarity between alleles *i* and *j* and *s*_*ii*_ = 0. The measure of similarity between two alleles of a microhaplotype used here was the reciprocal of the number of base pair differences between them.

These probabilities are then used to calculate the probability of observing each genotype given a true homozygous genotype. The probability of observing a genotype of BC given a true genotype of AA is

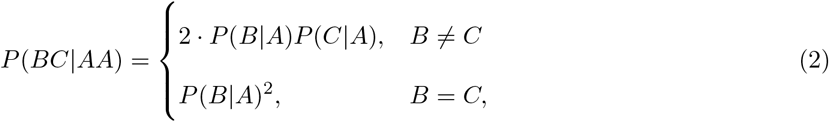

according to basic probability arguments, where *P* (*B*|*A*) is the probability of observing B when the true allele is A. The probability of observing each possible genotype given a true heterozygous genotype is calculated similarly, but also incorporates a probability of allelic dropout for each allele. Each allele is given a probability of dropping out, *d*_*i*_ for allele *i*, and in the current study a probability of 0.005 was used for all alleles. It is assumed that allelic dropout events are disjoint, and so the probability of no dropout occurring is 1 *− d*_*i*_ *− d*_*j*_. The probability of observing each genotype given no dropout can be calculated using the observation probabilities for each allele calculated in equation 1. The probability of observing a given genotype if a dropout has occurred is taken to be the probability of observing that genotype given a true homozygous genotype for the remaining allele. The total probability of observing a genotype is the sum of the probabilities of all three cases (no dropout, allele 1 dropout, allele 2 dropout).

